# High-throughput propagation of human prostate tissue from induced-pluripotent stem cells

**DOI:** 10.1101/637876

**Authors:** AC Hepburn, EL Curry, M Moad, RE Steele, OE Franco, L Wilson, P Singh, SE Crawford, Luke Gaughan, IG Mills, SW Hayward, CN Robson, R Heer

## Abstract

Primary culture of human prostate organoids is slow, inefficient and laborious. To overcome this, we demonstrate a new high-throughput model where rapidly proliferating and easily handled induced pluripotent stem cells, for the first time, enable generation of human prostate tissue in vivo and in vitro. Using a co-culture technique with urogenital sinus mesenchyme, we recapitulated the in situ prostate histology, including the stromal compartment and the full spectrum of epithelial differentiation. This approach overcomes major limitations in primary cultures of human prostate stem, luminal and neuroendocrine cells, as well as the stromal microenvironment. These models provide new opportunities to study prostate development, homeostasis and disease.

## Introduction

The capacity to generate in vitro 3D organoid cultures is transforming the study of human diseases (Clevers, 2016). These structures faithfully mimic in vivo epithelial architecture and present novel opportunities for pre-clinical studies (Gao et al., 2014; Karthaus et al., 2014). However, the widespread adoption of organoid culture in prostate studies is hampered by inherent shortcomings, including limited access to patient samples and the inefficient establishment of organoid cultures – limited to just the advanced metastatic cancers (Gao et al., 2014). In cases where longer term cultures are established, biological variability between patients and the genotypic and phenotypic drift through in vitro culture adaptation have hindered the utility of organoids in translational studies (Lee et al., 2018). Furthermore, current in vitro human prostate organoid approaches, from either tissue-derived cells or embryonic stem cells (ESCs), do not fully recapitulate the full breadth of in situ prostate differentiation as they do not contain neuroendocrine cells (Calderon-Gierszal and Prins, 2015; Karthaus et al., 2014). These shortcomings limit their utility, particularly imperative in light of emerging data on neuroendocrine differentiation driving treatment-resistant prostate cancer (Bishop et al., 2017). Although, human ESCs through co-engraftment with rodent urogenital sinus mesenchyme (UGM) can generate prostate tissue in vivo (Taylor et al., 2006), this ability was not demonstrated in vitro. Additionally, the use of alternative stem cell models would avoid significant ethical and regulatory restrictions relating to ESCs and also enable greater access to organoid generation to groups worldwide. Using induced pluripotent stem cells (iPSCs) to generate human prostate tissue would provide an easy-to-handle and rapidly proliferating source of cells delivering a solution to the problems of limited input from primary biopsies, ethics and access (Moad et al., 2013).

Herein, we demonstrate for the first time that tissue recombinants comprising human iPSCs and rat UGM generated both in vivo xenografts and in vitro prostate organoids that recreated the full breadth in situ prostate epithelial differentiation, including neuroendocrine cells, as well as the stromal compartment.

## Results

### Generation of human iPSC-derived prostate tissue in vivo

Firstly, as the tissue of origin used to generate the iPSCs can affect subsequent differentiation potential (Kim et al., 2011), we reprogrammed human prostate cells using a modified integration-free Sendai virus approach (Moad et al., 2013) (details in Supplementary Material and Methods). Reprogramming was confirmed by characteristic ESC morphology and marker expression, and importantly functional pluripotency in generating all three germ-layer lineages both in vitro and in vivo (Supplementary Fig. 1). Mimicking the in utero development of the prostate, which is driven by inductive UGM, we undertook sub-renal capsule grafting of iPSCs with UGM in nude mice (Cunha et al., 1987). This co-engraftment approach resulted in formation of prostatic tissue by 12 weeks (Fig. 1). Specifically, grafts resembled typical human prostate tissue histology, consisting mainly of glandular structures surrounded by myofibroblasts (Fig. 1a-b). UGM cells injected alone did not develop into glands, as also previously described (Taylor et al., 2006). The human origin of the epithelial cells was verified by immunolocalisation with anti-human mitochondria detection (Fig. 1c) and expression of cytokeratins CK8/CK18 on the cell surface and nuclear p63 demonstrated stratification of epithelium into characteristic prostate luminal and basal cells, respectively (Fig. 1d-e). Androgen receptor (AR) is an essential driver of prostate differentiation and both nuclear and cytoplasmic expression of the receptor was demonstrated (Fig. 1f). Furthermore, terminal differentiation was confirmed by the nuclear localisation in luminal epithelial cells of the prostate specific homeobox protein NKX3.1 and secretory prostate specific antigen (PSA) (Fig. 1g-h). Critically, Chromogranin A (ChrA) expression identified sparse neuroendocrine cells indicating that our bespoke methodology recapitulates the full breadth of prostate epithelial differentiation (Fig. 1i) (Blackwood et al., 2011).

**Fig. 1.**
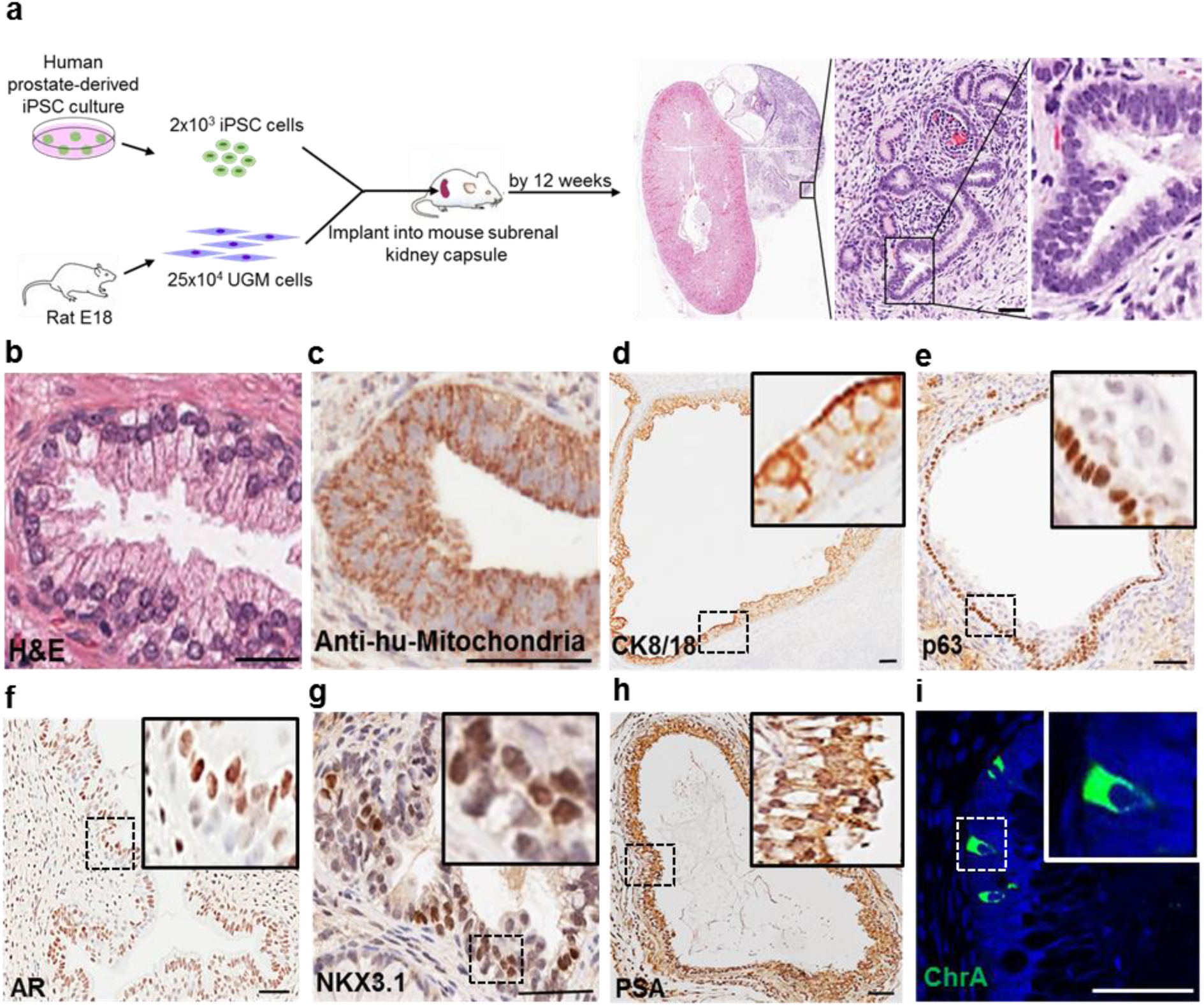
In vivo generation of human prostate tissue. (**a**) Schema overview. No invasion outside the renal capsule was seen consistent with benign histology. (**b**) Example of human prostate histology. (**c**) Xenograft showing anti-human mitochondria staining. (**d**) Luminal epithelium confirmed by CK8/18. (**e**) Basal epithelium confirmed by p63. (**f-h**) Luminal epithelium expressed AR, NKX3.1 and PSA. (**i**) Sporadic ChrA cells. Nuclei counterstained with DAPI. Scale=50μm. (n=12 mice)

### In vitro human iPSC-derived prostate organoids recapitulated the full breadth in situ prostate epithelial differentiation

Since UGM induced prostate differentiation from iPSCs in vivo, we hypothesised that it may also direct prostate differentiation in vitro and proceeded to employ a co-culture methodology (Fig. 2a). During embryogenesis, the prostate gland is derived from the endodermal urogenital sinus, thus directing differentiation of iPSCs down an endodermal lineage is likely to increase efficiency of prostatic epithelial differentiation. Accordingly, human prostate-derived iPSCs were differentiated through a definitive endoderm (DE) intermediary step (Supplementary Fig. 2) using Activin A and increasing concentrations of fetal calf serum (FCS) over 3 days, resulting in typical endodermal cobblestone-like morphology, increased cell size and reduction in the nuclear-to-cytoplasmic ratio (Supplementary Fig. 2a) (D’Amour et al., 2005). DE differentiation was confirmed by enrichment of DE-specific gene expression (SOX17 and FOXA2) (Supplementary Fig. 2b-c). Ten thousand of these cells were subsequently co-cultured with 35 × 10^3^ rat UGM cells in 3D matrigel culture (details in Supplementary Materials and Methods section). Formation of solid spherical structures was observed after 5 weeks that mimicked embryonic prostate organogenesis characterised by expression of the prostate-specific lineage marker NKX3.1 in cells with a basal phenotype (cytokeratin 34βE12 and p63) (Supplementary Fig. 3a-c). These structures occasionally contained small lumens associated with luminal marker expression (AR and CK8/18) (Supplementary Fig. 3d-e). Initial solid sphere formation primarily comprised of basal cells is known to proceed the generation of bi-layered organoids (basal and luminal layers) in both foetal prostate development and also in primary prostate organoid cultures (Karthaus et al., 2014; Xue et al., 1998).

**Fig. 2.**
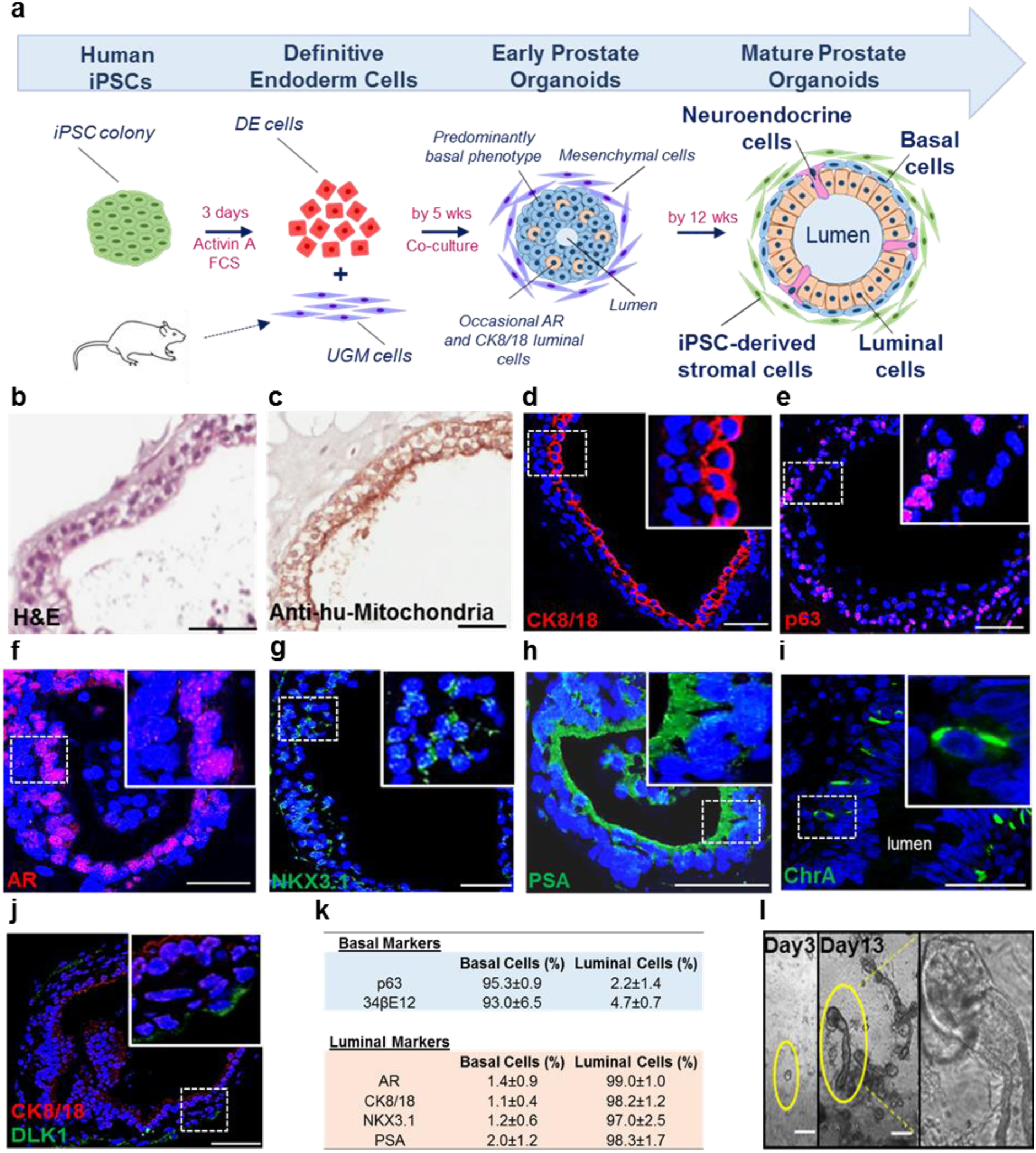
In vitro generation of human prostate organoids. **(a)** Schema overview. **(b)** Histology of organoids resembling prostate glands **(c)** Epithelium identified by anti-human mitochondria staining. **(d)** CK8/18 confirmed luminal cells. **(e)** p63 confirmed basal cells. **(f-h)** AR, NKX3.1 and PSA expression by luminal cells. **(i)** Rarely ChrA cells (4.4±1.3%, n=535 cells, n=3 experiments). **(j)** Subpopulation of basal cells expressed DLK1 (3.0±1.3%, n=650 cells, n=3 experiments). Scale=50μm. **(k**) Basal and luminal marker expression (n=3 experiments, n=183 organoids (164±33 cells/organoid)). **(l)** Day 3, early clone formation from 1-2 cells; Day 13, canalisation of tubular structures with dense bud tips. Scale=25μm.

**Fig. 3.**
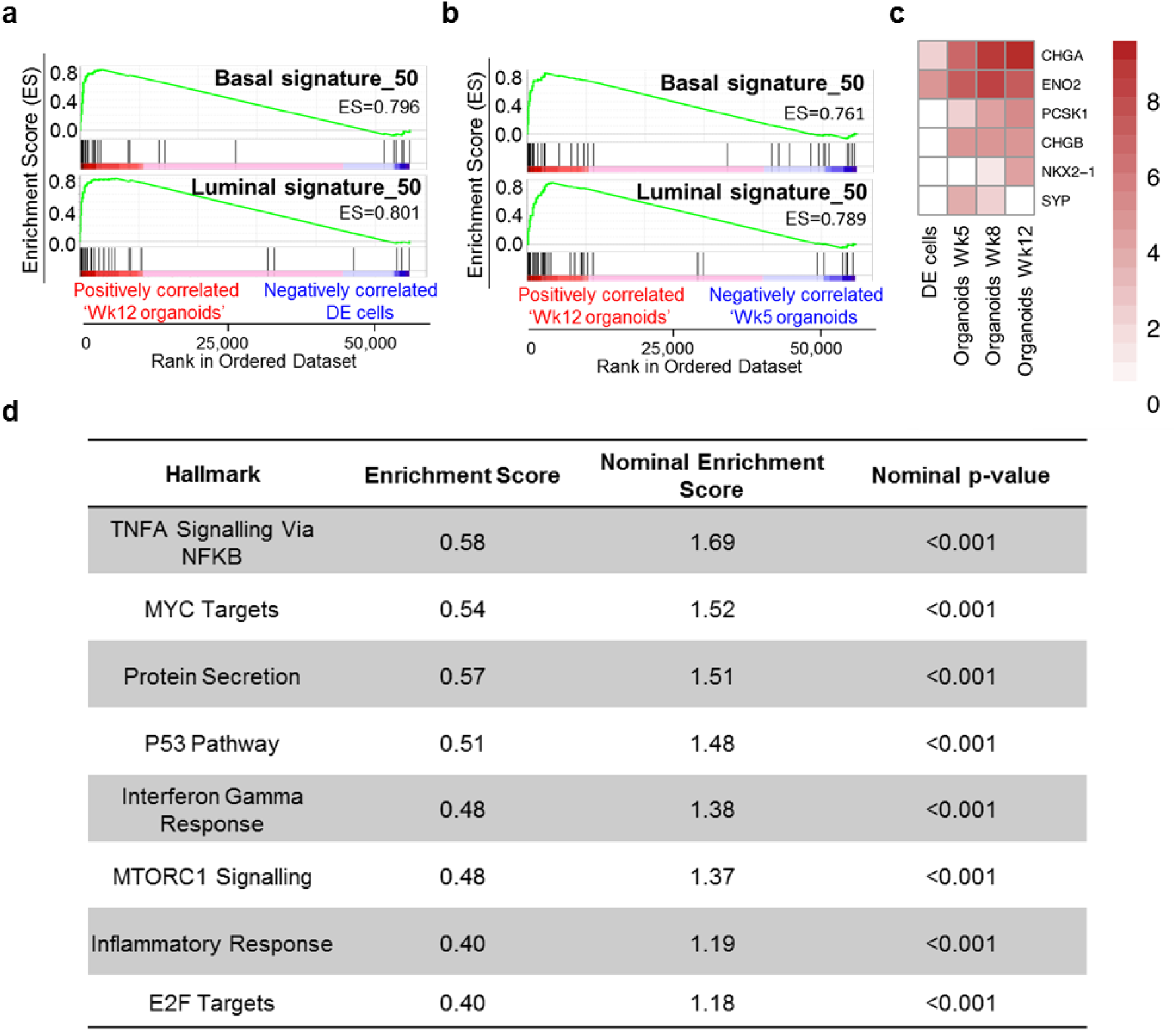
In vitro iPSC-derived prostate organoids shared gene expression profiles of mature human prostate cells. **(a)** GSEA demonstrating enrichment of basal and luminal gene expression in mature organoids (Wk12) in comparison to DE cells. **(c)** Same as in (b) but Wk12 vs Wk5 organoids. **(d)** Heatmap demonstrating neuroendocrine marker expression. Data is Log2 transformed. **(e)** GSEA Hallmark analysis identified 8 statistically significant enriched pathways in mature prostate organoids.

The ultimate objective was to replicate human prostate histology in vitro, by generating multi-layered prostate ductal-acinar epithelium composed of basal and luminal layers, and neuroendocrine cells within these solid spherical structures that can be visualised by expression of differentiation specific markers. To this end, prostate organoid culture medium (Karthaus et al., 2014), was applied to cultures from week 2 to support their growth and maintenance. These structures subsequently formed multi-layered acinar-like organoids with large lumens (size 65μm–455μm). By 12 weeks, they resembled human prostate tissue, demonstrating classical glands histology by forming an outer basal and inner luminal layer (Fig. 2a-b). These cells were identified as human by immunolocalization of anti-human mitochondria staining (Fig. 2c). Expression of CK8/18 and p63 appropriately localised to luminal and basal located epithelial cells respectively (Fig. 2d-e). Furthermore, luminal epithelial cells expressed nuclear AR and the terminally differentiated nature of the organoids was confirmed by NKX3.1 and secretory PSA (Fig. 2f-h). Sporadic neuroendocrine cells were again identified by chromogranin A (Fig. 2i). These data demonstrate that similar to the in vivo grafts, the in vitro organoids also faithfully recreated the full breadth of in situ prostate epithelial differentiation. Additionally, the somatic stem cell enrichment marker DLK1, which is known to mark cells essential for the long term maintenance of prostate epithelium, was expressed in a subpopulation of basal cells (3.0±1.3%, n=650 cells, n=3 organoid cultures) (Fig. 2j) (Moad et al., 2017). Spatially restricted expression of differentiation specific markers, stratified for basal (p63, 34βE12) and luminal (AR, CK8/18, NKX3.1 and PSA) organoid cells, was consistently confirmed (Fig. 2k). Also, following cellular disaggregation for passage beyond 12 weeks, 3D culture led to branching ductal structures (Fig. 2l), therefore fully recapitulating the human prostate epithelial histology.

### In vitro iPSC-derived prostate organoids shared gene expression profiles of mature human prostate cells

Having demonstrated the immunohistological presence of basal, luminal and neuroendocrine cells, we sought to undertake a broader transcriptomic characterisation of the degree of prostate specific differentiation. To dissect the specific ability of the iPSC-derived prostate organoids to recreate the two main epithelial cell compartments – the basal and luminal cells, we compared transcriptomes of iPSCs to primary basal (CD49f^+ve^) and luminal (CD26^+ve^) cells from freshly sorted whole human prostates (Goldstein et al., 2010; Karthaus et al., 2014; Moad et al., 2017) (Supplementary Fig. 4). Comparison of iPSCs to basal cells identified 1775 differentially expressed genes and comparison of iPSCs to luminal cells identified 1213 differentially expressed genes by at least 2-fold (adj. p value < 0.01). Basal and luminal gene-sets of the top 50 upregulated genes were derived from the differential gene expression analysis of iPSCs vs basal or luminal cells (Supplementary Tables S1-3). Gene Set Enrichment Analysis (GSEA) confirmed the mature organoids shared the terminally differentiated transcriptomic identities of the benchmark primary adult cells (Fig. 3a-b). Furthermore, enrichment of neuroendocrine marker expression was also seen in mature prostate organoids (Fig. 3c). Additionally, pathway ontology analyses revealed new insights into the mechanisms of differentiation, such as p53, inflammation and Myc related pathways (known key regulators in prostate cell differentiation and prostate carcinogenesis, providing focus for future studies (Fig. 3d) (Berger et al., 2014; Kwon et al., 2014).

**Fig. 4.**
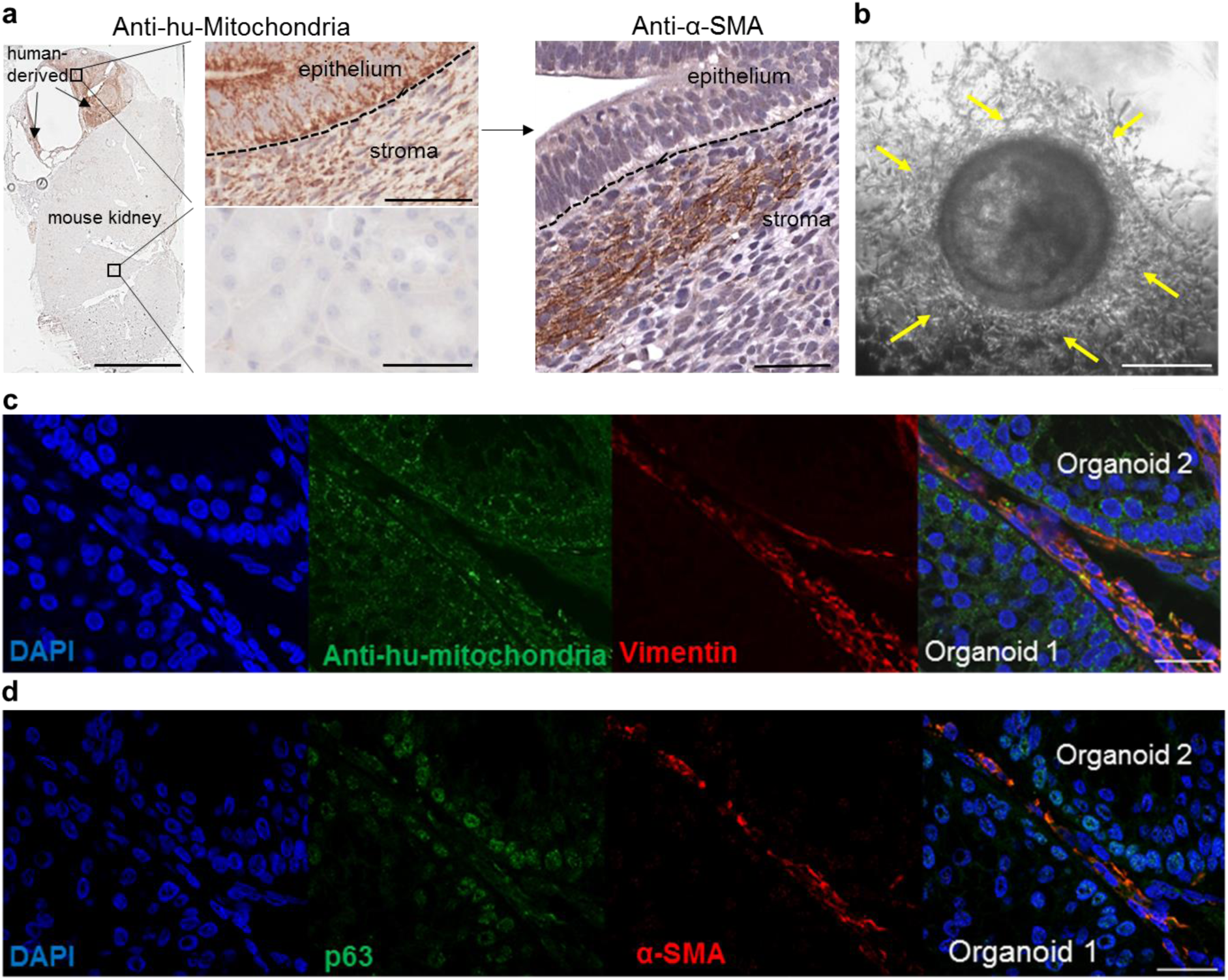
Prostate iPSCs generated a self-maintaining stromal compartment in mature prostate organoids. **(a)** Human anti-mitochondria and α-SMA co-localisation was seen in stromal cells within areas of the kidney containing the xenograft. Scale=50μm. **(b)** Stromal cells surrounded mature prostate organoids. Scale=25μm. **(c-d)** Overlapping expressions of vimentin, anti-human mitochondria, α-SMA and p63 demonstrated capacity of iPSCs to generate the stromal compartment. Scale=25μm.

### Prostate iPSCs generated a self-maintaining stromal compartment in mature prostate organoids

In the iPSC-derived prostate tissue xenografts, glands were surrounded by “nests” of cells with classical stromal morphology (Fig. 1a). These were demonstrated to be mesenchymal cells as shown by α-smooth muscle actin (α-SMA) expression and also of human origin as evidenced by expression analyses of the human specific anti-mitochondria marker (Fig. 4a). These data show that human-iPSCs also generated the stromal compartment. We next proceeded to examine whether the iPSCs demonstrated this ability in vitro. In the initial stages of differentiation, the UGM cells, whose inductive properties are transient as previously described (Xin et al., 2003), are ultimately replaced by human derived stromal cells, consistent with the reciprocal co-differentiation of both the epithelial and mesenchymal cells described in utero organogenesis provided the initial stromal compartment (Cunha et al., 1987). However, in mature in vitro-derived organoids, a self-maintaining stromal compartment derived from human iPSCs emerged (Fig. 4b). This was confirmed by α-smooth muscle actin (α-SMA) and vimentin co-localisation in anti-human mitochondria positive cells surrounding organoids (Fig. 4c-d).

## Discussion

In this report, we show for the first time that prostate iPSCs enabled generation of human prostate tissue both in vivo and in vitro. Using UGM-directed differentiation, human iPSC-derived prostate tissue comprehensively recapitulated in situ prostate histology and the full breadth of prostate specific epithelial differentiation – including neuroendocrine cells. During differentiation we saw the induction of master regulator transcription factors (AR and NKX3.1), which in mouse studies are central to prostate specific tissue programming (Talos et al., 2017). The archetypal histology of the human prostate is described by extensive acinar-tubular branching; however, previous organoid culture approaches have generated only acinar types of histology (Karthaus et al., 2014). Our work has shown that iPSC-derived organoid cultures also recapitulate this characteristic branching morphology, which is likely to be related to the stromal compartment (derived from the iPSCs) that is known to support branching (Richards et al., 2019).

Our approach provides new exciting opportunities to study and gain a more in-depth understanding of human prostate development, stem cell biology and homeostasis. Prostate cancer is the second commonest male cancer worldwide, accounting for 1.3 million new cases in 2018 (Bray et al., 2018). One of the key issues hindering the development of effective treatments for prostate cancer is the lack of suitable, tractable, and patient-specific in vitro models that accurately recapitulate this disease. High-throughput iPSC-derived human prostate tissue now provides unparalleled scope for in vitro disease modelling and drug discovery without the constraints of tissue accessibility and the long standing difficulties associated with primary culture. Here, in a complementary fashion to primary culture of prostate cancers, we propose a new concept where the generation of iPSC-derived normal tissue represents a malleable baseline model for genetic manipulation to recreate the disease of interest. Proof of concept is already established showing that benign prostate cells can be transformed into prostate cancers (Goldstein et al., 2010). Our proposed approach would overcome a major problem with the low efficiency of prostate cancer organoid culture and issues with significant genetic drift associated with long-term primary culture. Our baseline model, upon which mutations would be introduced, has the ability to reproduce, with high fidelity, isogenic cultures time after time.

The ability to differentiate iPSCs into human prostate tissue opens the door to unprecedented possibilities in the prostate research field. Freely shared, open access to these materials, together with the ease of the tissue generation using the simple protocol described, will result in an immediate impact for many researchers around the world.

## Materials and Methods

### Patient Material

All surgical specimens were collected according to local ethical and regulatory guidelines and included written, informed patient consent (Newcastle REC 2003/11 and Human Tissue Authority License 12534, Freeman Hospital, Newcastle Upon Tyne, UK).

### Human iPSC generation

Human prostate cell culture, characterisations by real time polymerase chain reaction, DNA fingerprinting, karyotyping, immunofluorescence, alkaline phosphatase staining and assays of pluripotency (embryoid body formation and teratoma formation) were described previously (Moad et al., 2013).

Pure cultures of 10 × 10^4^ prostate stromal cells seeded on twelve well plates were transduced using Cytotune 2.0 Sendai Virus reprogramming vectors (KOS, c-Myc and Klf4, Life Technologies) at a multiplicity of infection of 5-5-3 (KOS MOI=5, hc-Myc MOI=5, hKlf4 MOI=3) as recommended by the manufacturer’s instructions, in standard stroma culture medium (RPMI1640 medium with HEPES modification containing 10% foetal calf serum, 2mM L-glutamine and 1% penicillin and streptomycin (Sigma-Aldrich). On day 2, the transduction medium was replaced with fresh standard stroma culture medium. On day 7, cells were seeded onto vitronectin-coated twelve well plates at a concentration of 1.5 × 10^4^ cells per well in stroma culture medium. On day 8, medium was replaced with Essential 8 medium (Gibco) and changed every 48 h. From day 21, ESC-like colonies were manually selected based on morphology. Following clonal expansion and characterisation, iPSCs were cultured on hESC-qualified Matrigel (Corning) coated plates in mTeSR1 medium (Stem Cell Technologies). The medium was changed every 48 h.

### Definitive Endoderm Induction

To differentiate iPSCs into definitive endoderm (DE) cells, we modified the previously reported DE induction protocol (D’Amour et al., 2005), with other approaches also showing similar levels of efficiency (Loh et al., 2014). iPSCs were harvested to a single cell suspension using Gentle Cell Dissociation Reagent (StemCell Technologies) and plated at a density of 2× 10^6^ cells per well of Matrigel (Corning) coated six well plates in mTeSR1 medium (Stem Cell Technologies) containing 1μg/ml ROCK inhibitor Y-27632 (Stem Cell Technologies). Cells were incubated at 37°C for 24 h prior to incubation with DMEM/F12 medium (Sigma-Aldrich) supplemented with 100ng/ml Activin A (R&D Systems). Following 24 h, media was replaced with DMEM/F12 containing 100ng/ml Activin A and 0.2% foetal calf serum (FCS, Sigma-Aldrich) for a further 24 hours. Media was replaced with DMEM/F12 containing 100ng/ml Activin A and 2% FCS for a final 24h incubation.

### Tissue recombination grafts of human iPSCs with rat UGM

All animal experiments were performed in accordance with the Institutional Animal Care and Use Committee at North Shore University HealthSystem Research Institute, Evanston, IL. Pregnant Sprague-Dawley rats (Harlan Laboratories Inc, Indianapolis, IN, USA) were sacrificed at embryonic day 18. The embryos were isolated and urogenital systems removed. The UGS was separated from the bladder, urethra, Wolffian and Müllerian ducts and testes or ovaries and incubated in 10mg/ml trypsin (Sigma) at 4°C for 90 min followed by serial washes with RPMI-1640 (Sigma) supplemented with 10% FCS and 1% penicillin and streptomycin (Sigma). After separation from UGE, based on previously defined protocols for ESC-derived prostate differentiation (Taylor et al., 2006), 250,000 UGM cells, were resuspended with 2000 iPSCs in 40 μl of rat collagen matrix, plated as a plug and incubated at 37°C overnight in the presence of RPMI-1640 containing 10% FCS and 1% penicillin and streptomycin.

### Athymic nude mouse host xenografting

Male athymic nude mice (Hsd:Athymic Nude-Foxn1nu; Charles River Laboratories) aged 10 weeks were used for sub-renal capsule grafts. Following castration, a 1-cm skin incision along the dorsal midline was made, followed by another incision (approx. 6-8mm) of the body wall along the line of fat which runs parallel to the spine right above the kidney area. Then the kidney was exteriorised and a capsulotomy was made to prepare the subcapsular space for the grafts. Grafts were then placed underneath the renal capsule and maneuvered into various locations along the kidney. Two grafts were placed into each kidney (upper and lower poles) which was then reintroduced back into the mouse. Surgical incisions were closed with suture (body wall) and staples (skin). A testosterone pellet 25mg) was inserted s.c. into the scruff of the neck (Franco et al., 2011).

### Xenograft harvest and processing

Hosts were sacrificed from 6 weeks post grafting by anaesthetic (Penthobarbital) overdose followed by cervical dislocation. Grafts were harvested and kidneys removed en-bloc. Whole kidneys were placed in 10% neutral buffered formalin for 24 h. After fixation, kidneys were processed, paraffin embedded, and sections cut at 5 μm for staining and immunohistochemistry (IHC) (Franco et al., 2011).

### Co-culture of human iPSCs with rat UGM cells

For co-culture of UGM and DE cells, chamber slide wells were coated with 40μl of GFR-Matrigel (Corning) and set at 37°C for 20 minutes. 10,000 DE and 35,000 UGM cells were resuspended in GFR-Matrigel diluted with DMEM/F12 Ham (1:1, set at 37°C for 30 minutes in the chamber slide wells before addition of DMEM/F12 Ham containing 2% ITS (insulin, transferrin, selenium) (Gibco) and 10nM DHT (Sigma). After 7 days, media was changed to UGM conditioned media collected from whole pieces of UGM incubated in DMEM/F12 Ham containing 2% ITS and 10nM DHT as previously used for successful *in vitro* culture of UGS (Bryant et al., 2014). Medium was further supplemented with 1μg/ml ROCK inhibitor Y-27632 (Stem Cell Technologies) and replaced every 48h. From 2 weeks, the media was changed to prostate organoid medium (Karthaus et al., 2014). Wells were harvested from 6 weeks onwards for histology and RNA extraction. Wells for histology were removed as a Matrigel plug, fixed in 10% formalin overnight and processed before embedding into paraffin. For RNA extraction, the Matrigel was digested by incubation with dispase at 37°C until the gel was completely dissolved. The mixture was gently pipetted to further break up the Matrigel, and transferred to an Eppendorf for centrifugation at 2000rpm for 5 minutes. The supernatant was removed and the pellet snap frozen in isopentane and stored at −80°C.

### Immunohistochemistry

Immunohistochemistry was performed on FFPE sections (4 μm) that were initially deparaffinised and hydrated. Microwave antigen retrieval was carried out with citrate buffer pH6 to unmask surface antigens. Endogenous peroxidase activity was removed by blocking with 3% H2O2. Sections were then blocked in horse serum and incubated in primary antibody overnight 4^°^ C. The antibodies used were anti-human mitochondria (1:200, Abcam. This is a human specific antibody used in xenographic model research (Cho et al., 2017; Langer et al., 2018; Lee et al., 2017)), p63 (1:50, Leica Biosystems), CK8/18 (1:50, BD Pharmingen), AR (1:50, BD Pharmingen), NKX3.1 (1:50, AthenaES), PSA (1:25, Biogenex) and α-SMA (1:100, Abcam). Sections were washed and incubated with anti-rabbit or anti-mouse secondaries (ImmPRESS HRP Anti-Rabbit/Anti-Mouse IgG (Peroxidase) Polymer Detection Kit (Vector Laboratories). Antibody was detected with DAB solution (ImmPACT DAB Substrate Kit, Vector Laboratories) and counterstained with haematoxylin, dehydrated and mounted using DPX. Slides were then visualised using Aperio CS2 (Leica Biosystems).

### Immunofluorescence

Immunofluorescence was performed on frozen sections (4 μm). Sections were fixed with 4% paraformaldehyde, permeabilised using 0.1% triton and blocked in 4% BSA before incubation with primary antibody overnight at 4° C. The antibodies used were anti-human mitochondria (1:100, Abcam), p63 (1:50, Leica Biosystems), CK8/18 (1:50, BD Pharmingen), AR (1:50, BD Pharmingen), NKX3.1 (1:50, AthenaES), PSA (1:50, Biogenex), Vimentin (1:100, Abcam), α-SMA (1:100, Abcam), FOXA2 (1:10, R&D) and SOX17 (1:20, R&D). Secondary antibodies Alexa Fluor-488 (1:100; ab150129; Abcam, UK), Alexa Fluor-568 (1:100; A-10042; Invitrogen, USA), Alexa Fluor-546 (1:100; A-11030; Life Technologies, USA) were used to detect the bound unconjugated primary antibody. Sections were washed and mounted using Vectashield with DAPI mountant (Vector Laboratories, Peterborough, UK) before being visualised on Leica SPE confocal and widefield fluorescence inverted microscope (Leica Biosystems).

### RNA extraction, reverse transcription and real time PCR (qPCR)

Total RNA was extracted using Ribozol™ RNA Extraction Reagent (Amresco) and reverse transcribed using Moloney murine leukaemia virus reverse transcriptase enzyme (Promega) according to the manufacturer’s instructions. Real time PCR (qPCR) was carried out using Platinum SYBR® green qPCR SuperMix-UDG (Invitrogen) in 384-well clear optical reaction plates using the ABI 7900HT qPCR system (Applied Biosystems) according to the manufacturer’s instructions. Levels of expression were normalised against housekeeping gene GAPDH. Primers were: FOXA2 F: 5’-TCCGACTGGAGCAGCTACTATG-3’ and R: 5’-CCACGTACGACGACATGTTC-3’; GAPDH F: 5’-CGACCACTTTGTCAAGCTCA-3’ and R: 5’-GGGTCTTACTCCTTGGAGGC-3’.

### RNA sequencing analysis

Total RNA was extracted from cells using Ribozol^™^ RNA Extraction Reagent (Amresco) following manufacturer’s instructions. RNA-Seq library construction and sequencing was performed at Otogenetics Corporation (Atlanta, USA) according to standard protocols. Resulting RNA-Seq fastq reads were aligned to Hg19 (GRCh37) using STAR (Dobin et al., 2013) and mapped to genes using HTSeq counts (http://htseq.readthedocs.io/en/master/count.html). Normalised count and differential expression analysis data was generated using DESeq2 (Love et al., 2014). Gene Set Enrichment Analysis (GSEA) (Mootha et al., 2003; Subramanian et al., 2005) was performed on normalised RNA-seq count data and calculated by permuting genes 1000 times in the GSEA software. All heatmaps were generated using R3.4.2.

## Acknowledgements

Funding from Cancer Research UK, Medical Research Council and Prostate Cancer UK.

