## Supplementary Data for "High-throughput propagation of human prostate tissue from induced-pluripotent stem cells"

**Supplementary Figure 1**

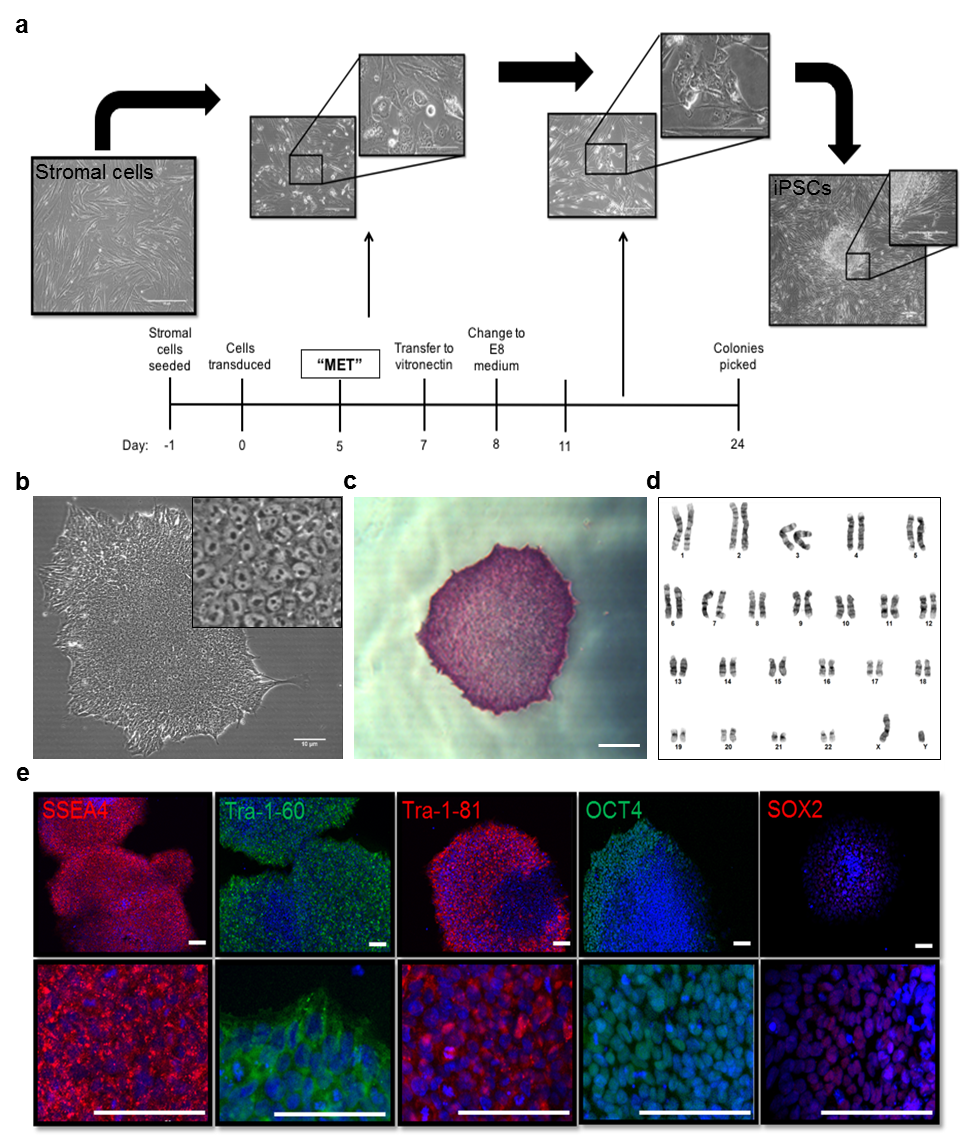

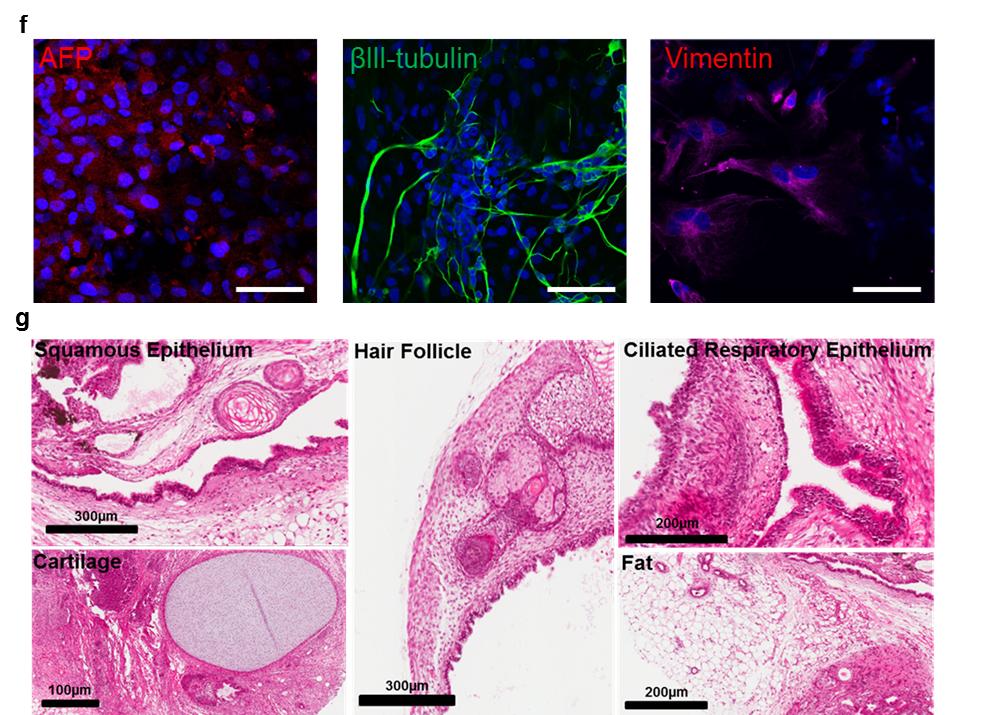

**Supplementary Figure 1. Reprogramming human prostate cells into iPSCs.**

**(a)** We have previously shown that tissue from prostate mesoendodermal lineage is able to generate prostate specific differentiation using an integrative polycistronic lentiviral vector (Moad et al., 2013). Here we show a schematic of the timescale for reprogramming primary prostate fibroblasts to iPSCs using integration free Cytotune 2.0 Sendai viral vectors. Micrographs show the change in cell morphology over this period from mesenchymal-epithelial transition (MET) to appearance of ESC-like colonies (n=3 clones). **(b)** ESC-like morphology of prostate iPSC colony cultured using feeder-free conditions. Inset, magnified view. Scale bar 10 μm. **(c)** Alkaline phosphatase staining of prostate iPSC colony. Scale bar 10 μm. **(d)** Prostate iPSCs confirmed to possess a diploid 46XY karyotype. **(e)** Immunofluorescence of prostate iPSCs for the expression of specific human ESC surface markers: stage specific embryonic antigen-4 (SSEA4), tumour rejection antigen (TRA)-1-60, TRA-1-81, and nuclear transcription factors OCT4 and SOX2. Bottom panel, magnified view. Scale bars 100 μm. **(f)** Immunofluorescence analysis of embryoid bodies derived from prostate iPSCs showing expression of the lineage markers α-fetoprotein (AFP, endodermal marker, left panel), βIII-tubulin (ectodermal marker, middle panel) and vimentin (mesodermal marker, right panel). Scale bars 25 μm. Nuclei were counterstained with 4′,6-diamidino-2-phenylindole (blue). **(g)** Histologic sections of teratoma formed from prostate iPSCs representing all three embryonic germ layers. Scale bars 100, 200 and 300 μm.

**Supplementary Figure 2**

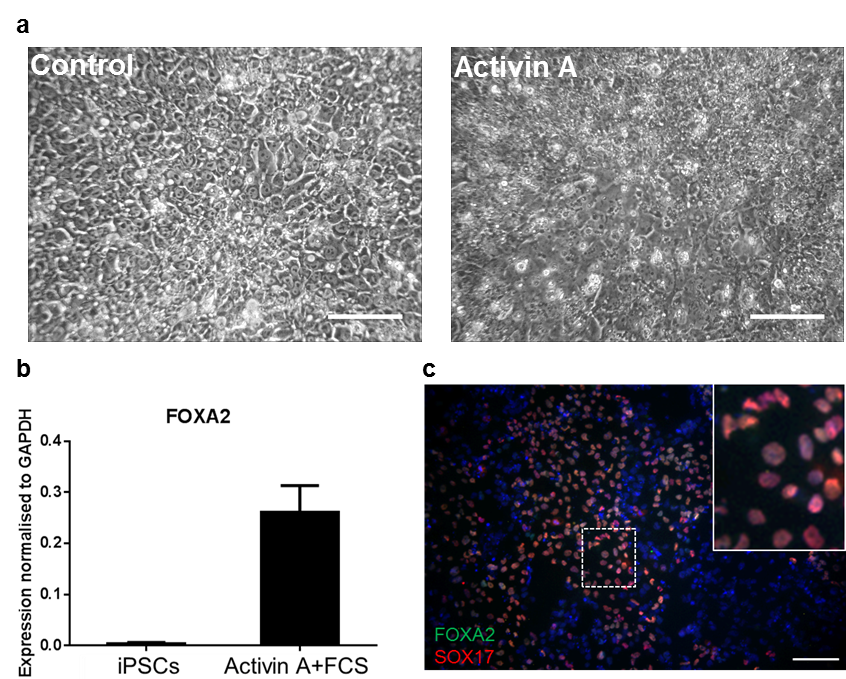

**Supplementary Figure 2: Formation of definitive endoderm from iPSCs.**

**(a)** Morphological changes of iPSCs at 72h following treatment with Activin A and FCS compared to control (untreated iPSCs). (**b**) Real time PCR analysis demonstrating expression of definitive endoderm (DE) specific marker FOXA2 following induction of prostate iPSCs with Activin A and FCS for 72h (Data represents at least three independent experiments ± SEM). **(b)** Immunofluorescence analysis demonstrating expression of DE specific markers FOXA2 and SOX17 following treatment of iPSCs with Activin A and FCS for 72h. Efficiency of DE differentiation was approximately 80%. Inset, magnified view. Scale bar 10 μm. (n=3 iPSC clones, n=3 assays per clone).

**Supplementary Figure 3**

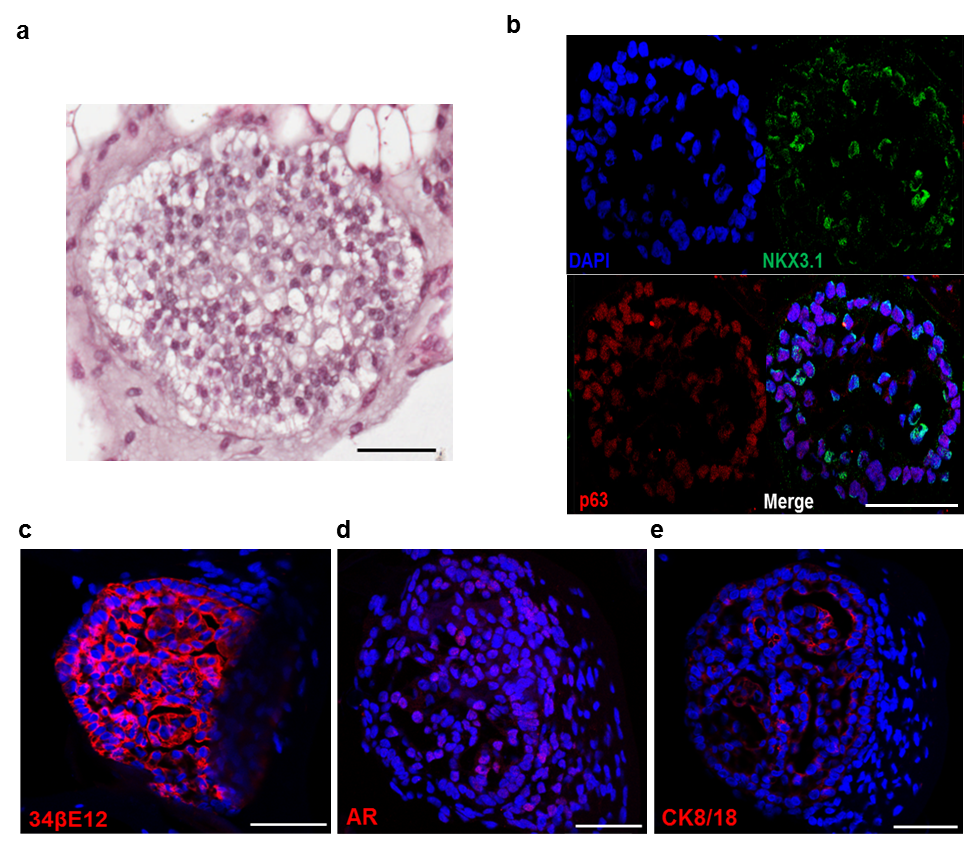

**Supplementary Figure 3: Characterisation of early prostate organoids.**

**(a)** Histology of early organoids demonstrating solid spherical structures. **(b)** Early organoids also predominantly expressed basal marker p63 and luminal transcription factor NKX3.1. **(c)** Predominant expression of basal cytokeratin 34βE12, which was expressed almost uniformly throughout. **(d)** Occasional expression of the transcription factor AR. **(e)** Sparse, expression of luminal cytokeratin CK8/18 was also seen. Scale bars 50 μm.

Although infrequently areas of early lumen formation were noted, on the whole we saw amorphous early spheroids consistent with reports of mixed luminal and basal phenotypes before clear differentiation into mature luminal and basal cell histology and restricted expressions of associated differentiation marks. These findings are consistent with the early stages of human fetal prostate development, with an hierarchical pathway of cellular differentiation from basal to luminal cells (Bhatia-Gaur et al., 1999, Xue et al., 1998).

**Supplementary Figure 4**

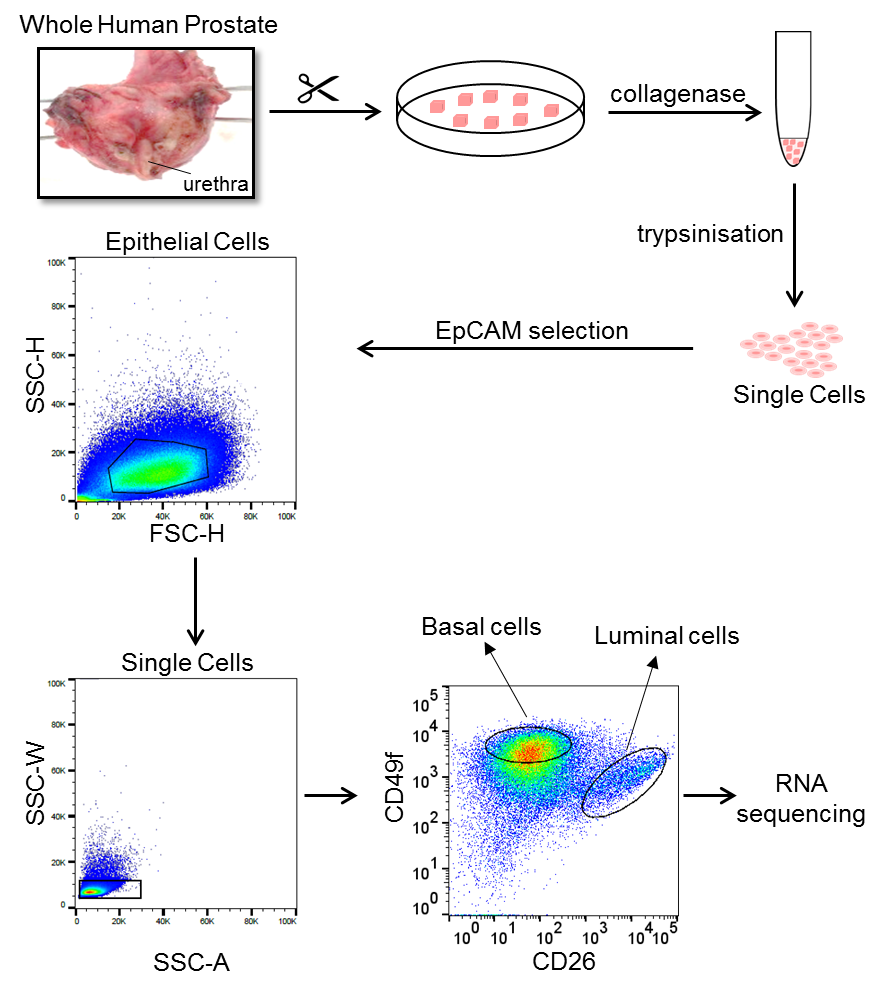

**Supplementary Figure 4: Strategy to isolate basal and luminal cells from whole human prostate.**

Whole human clinically benign prostates (n=3) from patients undergoing radical prostatectomy (catheter in urethra) for bladder cancer were processed to isolate basal and luminal epithelial cells for RNA sequencing (as previously described, Moad et al., 2017). Briefly, tissue was cut into small chunks and incubated with collagenase. Following trypsinisation, single epithelial cells were further enriched by performing MACS EpCAM selection. Samples were stained with basal CD49f and luminal CD26 markers before being FACS sorted. Size gating was applied to enrich for whole cells and doublet discrimination was undertaken to avoid false positive measures. Circular gates identify CD49f^+ve^ basal and CD26^+ve^ luminal cells based on isotype controls.

**Download spreadsheets for:**

**Supplementary Table S1.** Differential Gene Expression of iPSCs vs CD49f positive basal cells isolated from whole human prostates by flow cytometry (related to Fig. 3 and Supplementary Fig. 4). A list of the genes from RNA sequencing (n=3).

**Supplementary Table S2.** Differential Gene Expression of iPSCs vs CD26 positive luminal cells isolated from whole human prostates by flow cytometry (related to Fig. 3 and Supplementary Fig. 4). A list of the genes from RNA sequencing (n=3).

**Supplementary Table S3.** List of the top 50 genes upregulated in the DEG analysis to generate ‘basal’ and ‘luminal’ genesets for GSEA.

**Supplementary Data References**

Bhatia-Gaur, R., Donjacour, A.A., Sciavolino, P.J., Kim, M., Desai, N., Young, P., Norton, C.R., Gridley, T., Cardiff, R.D., Cunha, G.R., et al. (1999). Roles for Nkx3.1 in prostate development and cancer. Genes Dev 13, 966-977.

Moad, M., Pal, D., Hepburn, A.C., Williamson, S.C., Wilson, L., Lako, M., Armstrong, L., Hayward, S.W., Franco, O.E., Cates, J.M., et al. (2013). A novel model of urinary tract differentiation, tissue regeneration, and disease: reprogramming human prostate and bladder cells into induced pluripotent stem cells. Eur Urol 64, 753-761.

Moad, M., Hannezo, E., Buczacki, S.J., Wilson, L., El-Sherif, A., Sims, D., Pickard, R., Wright, N.A., Williamson, S.C., Turnbull, D.M., et al. (2017). Multipotent Basal Stem Cells, Maintained in Localized Proximal Niches, Support Directed Long-Ranging Epithelial Flows in Human Prostates. Cell Rep 20, 1609-1622.

Xue, Y., Smedts, F., Debruyne, F.M., de la Rosette, J.J., and Schalken, J.A. (1998). Identification of intermediate cell types by keratin expression in the developing human prostate. Prostate 34, 292-301.
